# Shared and divergent phase separation and aggregation properties of brain-expressed ubiquilins

**DOI:** 10.1101/2020.08.12.248401

**Authors:** Julia E. Gerson, Hunter Linton, Jiazheng Xing, Alexandra B. Sutter, Fayth S. Kakos, Jaimie Ryou, Nyjerus Liggans, Lisa M. Sharkey, Nathaniel Safren, Henry L. Paulson, Magdalena I. Ivanova

**Affiliations:** Department of Neurology, University of Michigan, Ann Arbor, MI 48109-2200; Neuroscience Graduate Program, Northwestern University Feinberg School of Medicine, Chicago, IL 60611, USA; Biophysics Program, University of Michigan, Ann Arbor, MI 48109-2200, USA

**Keywords:** proteostasis, ubiquilin, protein aggregation, neurodegenerative disease, phase separation, amyotrophic lateral sclerosis, frontotemporal dementia, fibril, FRAP

## Abstract

The brain-expressed ubiquilins, UBQLNs 1, 2 and 4, are highly homologous proteins that participate in multiple aspects of protein homeostasis and are implicated in neurodegenerative diseases. Studies have established that UBQLN2 forms liquid-like condensates and accumulates in pathogenic aggregates, much like other proteins linked to neurodegenerative diseases. However, the relative condensate and aggregate formation of the three brain-expressed ubiquilins is unknown.

We report that the three ubiquilins differ in aggregation propensity, revealed by *in-vitro* experiments, cellular models, and analysis of human brain tissue. UBQLN4 displays heightened aggregation propensity over the other ubiquilins and, like amyloids, UBQLN4 forms ThioflavinT-positive fibrils *in vitro*. Measuring fluorescence recovery after photobleaching (FRAP) of puncta in cells, we report that all three ubiquilins undergo liquid-liquid phase transition. UBQLN2 and 4 exhibit slower recovery than UBQLN1, suggesting the condensates formed by the brain-expressed ubiquilins have different compositions and undergo distinct internal rearrangements.

We conclude that while all brain-expressed ubiquilins exhibit self-association behavior manifesting as condensates, they follow distinct courses of phase-separation and levels of aggregation. This variability among ubiquilins along the continuum from liquid-like to solid likely informs both the normal ubiquitin-linked functions of ubiquilins and their accumulation and potential contribution to toxicity in various neurodegenerative diseases.

## Introduction

The ubiquilins are highly homologous proteins that regulate multiple pathways of protein homeostasis. Of the mammalian ubiquilins, UBQLN1, UBQLN2 and UBQLN4 are expressed in the brain and are associated to varying degrees with neurodegenerative diseases characterized by protein misfolding, aggregation and mislocalization. Mutations in UBQLN2 cause a rare familial X-linked neurodegenerative disease belonging to the amyotrophic lateral sclerosis/frontotemporal dementia (ALS/FTD) spectrum ^1,2^. Both UBQLN1 and UBQLN2 colocalize with disease aggregates in ALS/FTD, Huntington’s disease, and to a lesser extent, the synucleinopathies ^3,4^. A genetic variant of UBQLN4 has also been implicated in ALS, suggesting that UBQLN4, like UBQLN1 and UBQLN2, contributes to neurodegeneration ^5^.

The involvement of the ubiquilin family in protein accumulation associated with neurodegenerative disease likely reflects their functions in protein quality control. All three brain-expressed ubiquilins contain a C-terminal ubiquitin-associated (UBA) domain, an N-terminal ubiquitin-like (UBL) domain and four stress-induced protein-like (STI) domains that mediate binding to molecular chaperones. The most well-characterized function for ubiquilins is as ubiquitin-proteasome shuttle factors ^6,7^, but they have been implicated in other protein homeostasis pathways including autophagy and endoplasmic reticulum-associated protein degradation (ERAD) ^8–12^.

Like many other proteins associated with neurodegenerative diseases ^13–18^, UBQLN2 spontaneously phase separates to form condensates, or liquid droplets, in which proteins are concentrated yet remain mobile ^13,19,20^. Liquid-like condensates are tightly regulated by cells and can provide sites for specific cellular functions ^21–24^. Evidence for this includes a recent finding that UBQLN2 regulates the fluidity of protein– RNA complexes in FUS-related ALS/FTD ^25^. At the same time, dysregulated condensate formation by various proteins leads to abnormal protein accumulation and disease ^26,27^. *In vitro*, high salt concentrations promote UBQLN2 phase separation ^19,20^. Large punctate UBQLN2 assemblies resembling liquid condensates form in cells overexpressing UBQLN2 ^13^. In particular, mutations in a proline-rich domain present in UBQLN2 mediate the propensity of UBQLN2 to form less mobile liquid condensates and fibrillar aggregates ^13,20^, suggesting that dynamic changes to ubiquilin condensation may underlie toxicity in mutation-driven disease.

The liquid phase transition behavior and aggregation propensity of the other ubiquilins remain unknown. Because UBQLN1 and UBQLN2 form or co-localize with pathological deposits in disease ^3,28^, and all three brain-expressed ubiquilins are highly homologous proteins ^29^, we sought to define their relative phase separation and aggregation properties. Through *in vitro* studies, cellular models, and analyses of human tissue, we show that the three brain-expressed ubiquilins differ in condensate formation and aggregation. The results support the view that these highly homologous proteins exhibit complex association behavior and suggest that an imbalance in phase transitions may play a role in neurotoxicity related to ubiquilin protein dysregulation.

## Results

### Among the brain-expressed ubiquilins, UBQLN4 is particularly prone to self-assemble into fibrillar aggregates *in vitro*

We previously identified the UBA domain as a key driver of UBQLN2 aggregation ^13^. The UBA domains of all three brain-expressed ubiquilins are highly conserved, sharing 93% sequence identity (Fig. 1a) ^29^. This similarity led us to investigate whether, like UBQLN2, UBQLN1 and UBQLN4 can also aggregate. Using a predictor algorithm of amyloid-like structures ^30^, we calculated the aggregation propensity profile of the three constructs used in our *in-vitro* studies (Fig. 1b). These constructs (UBQLN1^438-589^, UBQLN2^430-624^ and UBQLN4^444-601^) extend from the fourth STI1 motif to the end of the protein (Fig. 1). Whereas the UBA domains have nearly identical aggregation profiles, the fibril formation profiles differ for the three constructs in the region between the fourth STI1 motif and the UBA domain (Fig. 1b). This region (fourth STI1 to UBA) also has the largest sequence variability across the ubiquilin constructs used for the *in vitro* studies and, in UBQLN2, is distinguished by its unique proline-rich (PXX) domain.

**Fig. 1.**
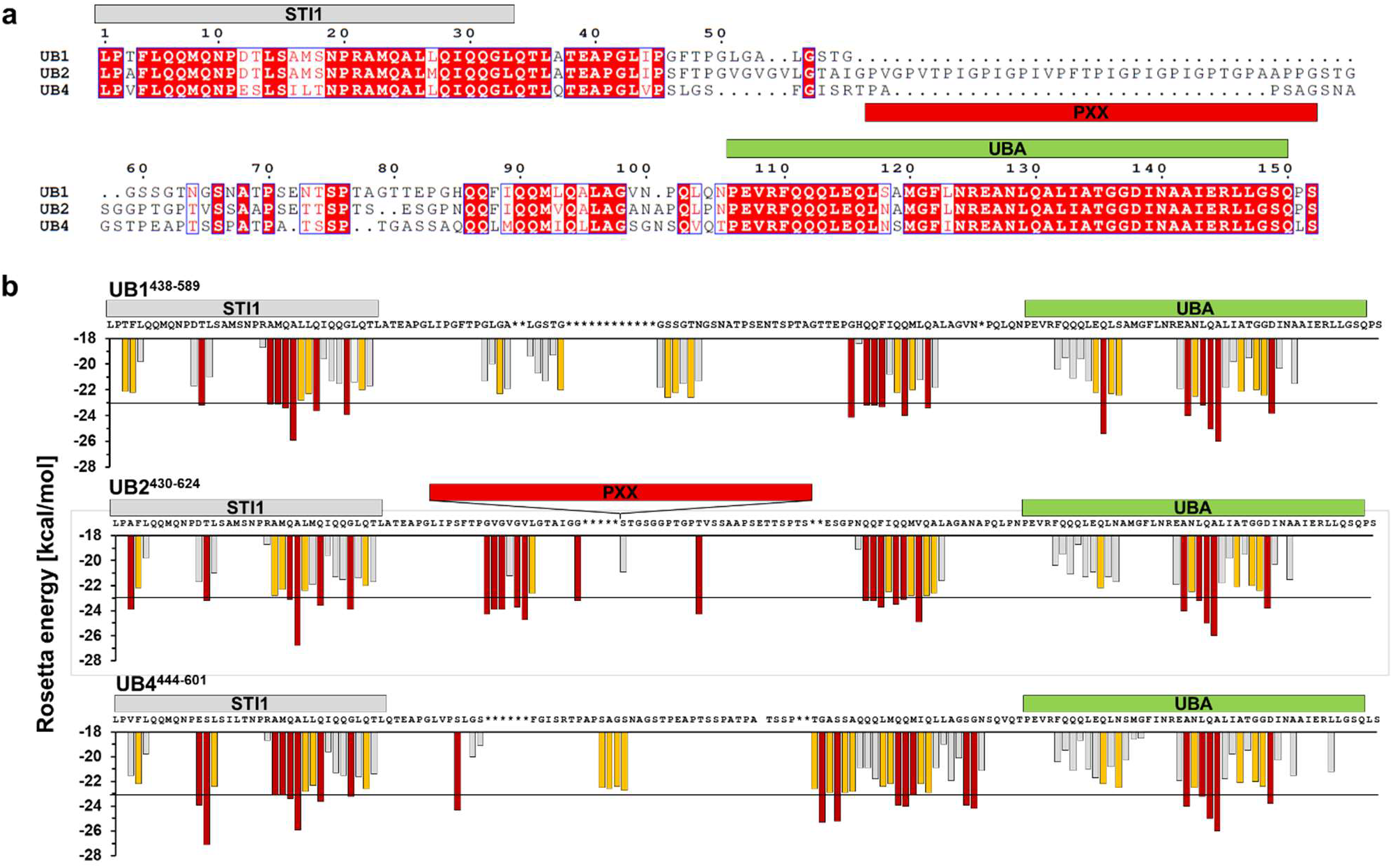
Sequence alignment and prediction of aggregation-prone regions of C-terminal ubiquilin constructs (UBQLN1^438-589^, UBQLN2^430-624^, and UBQLN4^444-601^) used for *in vitro* studies. **(a)** The UBA domains are highly conserved (93% identical) with the major sequence differences residing in the region between STI1-4 and UBA ^29,48^. Identical residues are colored in red. **(b)** Zipper DB was used to predict regions that may drive aggregation of ubiquilin constructs ^30^. In the histogram, each bar represents a six-residue-long segment. Segments with Rosetta energies smaller than −23 kcal/mol (black line) are predicted to form β-sheet structures characteristic of amyloids, and are colored in red. In orange are the segments where Rosetta energies are between −22 and −23 kcal/mol, a unit larger than the threshold of −23 kcal/mol. As with the sequence alignment, the aggregation profiles of these ubiquilin constructs differ mainly in the region between STI1-4 and the UBA domain. The PXX repeat motif of UBQLN2 is not shown because none of the segments were predicted to form β-sheet structures. Asterisks designate the gaps in the multiple sequence alignment of the ubiquilins.

Because the UBA domain is known to be critical for UBQLN2 aggregation and phase separation ^13,19^, we anticipated that recombinant UBQLN2^430-624^ and UBQLN4^444-601^ would similarly aggregate *in vitro.* To measure the aggregation, we used ThioflavinT (ThT), a dye that produces characteristic fluorescence in the presence of β-sheet structures and has been used to study fibril formation of many amyloid proteins^31,32^.

Among the three ubiquilins, UBQLN4^444-601^ aggregated most rapidly (Fig. 2). At a 10 μM concentration, the aggregation reaction of UBQLN4^444-601^ was the fastest to transition to a steady-state, characterized by a constant fluorescence over time. Indeed, the time to half transition to steady-state for UBQLN4^444-601^ was similar to that of the isolated UBA domain from UBQLN2, which drives aggregation of the full protein ^13,19^. In contrast, UBQLN1^438-589^ and UBQLN2^430-624^ aggregated more slowly and displayed decreased ThT signal (Fig. 2b).

**Figure 2.**
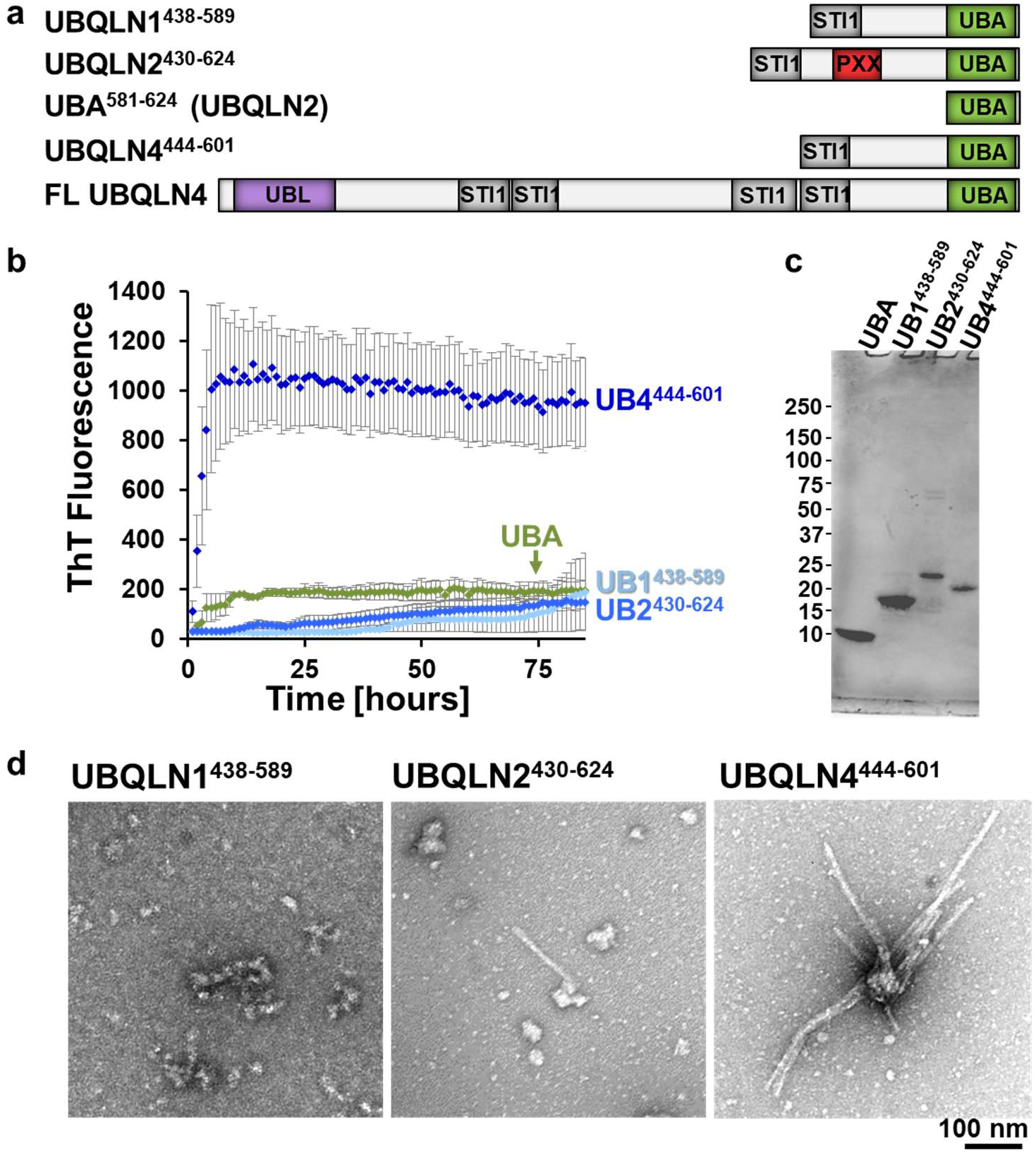
UBQLN4^444-601^ displays greater propensity to form amyloid-like aggregates than UBQLN1^438-589^ or UBQLN2^430-624^. **(a)** Constructs used for the studies of protein aggregation *in vitro*. The UBA^581-624^, a known driver of UBQLN2 aggregation, was tested as a positive control. FL UBQLN4 is shown for comparison on the bottom. **(b)** ThT assay was used to monitor aggregation of ubiquilin constructs. The fluorescence is the mean of four or more independent experiments (bars =SEM). **(c)** SDS-PAGE of the recombinant proteins used in the ThT assays. **(d)** Samples for TEM imaging were collected at the termination of ThT assays. UBQLN4^**444-601**^ samples produced fibrillar species resembling amyloid fibrils and UBQLN2^**430-624**^ formed a heterogeneous mixture of small fibrils and globular species, while no fibrils were observed for UBQLN1^**438-589**^.

We note that for this study we used a UBA construct (UBA^581-624^) that is seven residues shorter than UBA^574-624^, which we previously described ^13^. UBA^581-624^ also lacks the thrombin cleavage site and a linker sequence present in the longer UBA^574-624^ construct ^13^. These differences likely explain the shortened lag phase of UBA^581-624^ compared to UBA^574-624^ ^13^.

At reaction completion, aggregated samples were viewed by transmission electron microscopy (TEM). UBQLN4^444-601^ readily formed fibrillar aggregates resembling amyloid fibrils. In contrast, UBQLN1^438-589^ and UBQLN2^430-624^ mainly formed spherical species, though samples of UBQLN2^430-624^ had low levels of sparse, short fibrils (Fig. 2d). The formation of amyloid-like fibrils by UBQLN4^444-601^ likely explains its increased ThT fluorescence at steady-state, as ThT binds β-sheet structures characteristic of amyloids.

### Among tested ubiquilins, UBQLN4 is most prone to partition into the PBS-insoluble fraction of cell lysates

In our *in vitro* ThT assays we used truncated ubiquilin constructs that included the region of the protein described to drive phase separation and aggregation ^13,19^. To determine whether the divergent aggregation behavior of ubiquilins observed *in vitro* extended to cells, we chose to analyze full-length proteins. We transiently expressed full-length FLAG-tagged UBQLNs 1, 2 and 4 (FLAG-UBQLN1, FLAG-UBQLN2, and FLAG-UBQLN4) in HEK-293 cells and separated cell lysates into PBS-soluble (supernatant) and PBS-insoluble fractions (pellet). Because our lysis protocol using mechanical shearing is efficient in releasing soluble proteins that normally reside in the nucleus (Supplemental Fig. S1), the majority of the proteins in the PBS-insoluble fraction are likely to be in an aggregated state as opposed to trapped in a subcellular structure. All three proteins partitioned into both PBS-soluble and insoluble fractions. Compared to FLAG-UBQLN1 and FLAG-UBQLN2, a significantly higher percentage of FLAG-UBQLN4 partitioned into the PBS-insoluble fraction (Fig. 3a-b). Moreover, only FLAG-UBQLN4 was noted to electrophorese as a high molecular weight (HMW) smear on denaturing Western blots (Fig. 3c). This pattern of relative solubility/insolubility and formation of SDS-resistant HMW species by full-length ubiquilins mirrors the relative aggregation behavior of the C-terminal ubiquilin constructs *in vitro*.

**Figure 3.**
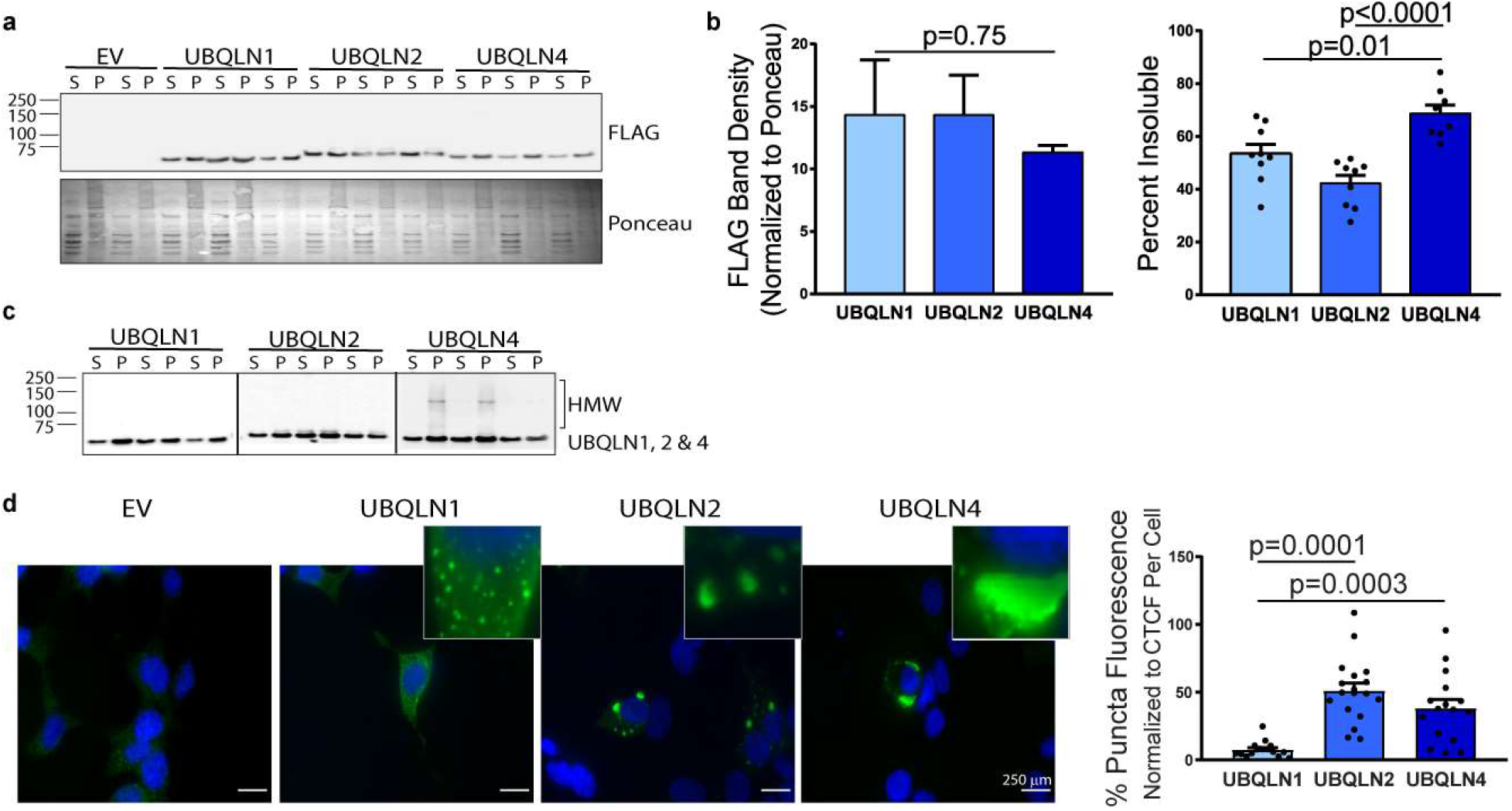
In transfected cells, UBQLN4 and UBQLN2 display enhanced aggregation and puncta formation compared to UBQLN1. **(a)** Of the three tested ubiquilins expressed in HEK-293 cells, FLAG-UBQLN4 is most prone to deposit in PBS-insoluble fractions (n=9). **(b)** All three ubiquilins were expressed equally as measured by total FLAG band density (normalized to Ponceau). For measurements of insolubility, values are expressed as percentage of protein that is PBS-insoluble, determined from the calculated total pool of PBS-soluble (S) and PBS-insoluble (P). **(c)** UBQLN1, 2 and 4-specific antibodies (cropped to present all three antibodies in parallel) show that UBQLN4 partially electrophoreses as detergent-resistant high molecular weight species whereas UBQLNs 1 and 2 do not (uncropped Western blots are shown in Supplemental Figure S2). **(d)** Immunofluorescence of FLAG-UBQLNs 1, 2 and 4 shows that FLAG-UBQLN1 in HEK-293 cells is primarily diffusely expressed in the cytoplasm, and only a fraction of the protein is localized in small, spherical puncta. In contrast, a higher percentage of FLAG-UBQLNs 2 and 4 are sequestered into large, heterogeneous puncta when normalized to corrected total cell fluorescence (CTCF) (n= 20; right plot). Data were analyzed by one-way ANOVA using the Bonferroni post-hoc test. Means and SEMs are displayed. Scale bar = 250 μm.

To further evaluate assembly formation and aggregation of each ubiquilin, we quantified the formation of intracellular puncta by each protein. Assessed by immunofluorescence, FLAG-UBQLN1 primarily distributed diffusely in cells whereas a much greater percentage of FLAG-UBQLN2 and FLAG-UBQLN4 localized to puncta (Fig. 3d).

### Divergent ubiquilin solubility in human brain

To gain insight into the physiological relevance of this divergence in ubiquilin aggregation, we analyzed levels of each brain-expressed ubiquilin in the PBS-soluble versus PBS-insoluble fractions of brain lysates derived from human brain tissue, assessing both normal controls and various neurodegenerative diseases.

Results in human brain samples supported our results *in vitro* and in cellular models: the percentage of UBQLN4 present in the PBS-insoluble fraction was elevated over that of UBQLN1. The relative levels of UBQLN2 insolubility varied between samples but appeared to be intermediate between UBQLN1 and UBQLN4 (Fig. 4). We assessed cingulate cortex from two synucleinopathies, Parkinson’s disease (PD) and dementia with Lewy bodies (DLB), and frontal cortex from the tauopathy progressive supranuclear palsy (PSP), as well as cortex from age-matched control brains (Supplemental Table S1). Comparisons of the soluble and insoluble fractions from both PD and DLB demonstrated that the behavior of UBQLN2 and UBQLN4 are similar, partitioning into the insoluble fraction to a greater extent than UBQLN1. Similarly, in PSP brain tissue, UBQLN4 insolubility was significantly elevated over UBQLN1 and UBQLN2. The inherent heterogeneity of human disease tissue samples likely explains the sample-to-sample variability in degree of solubility for the three ubiquilins, but UBQLN4 displayed the least variation, consistently showing a high degree of insolubility in all three diseases, as well as in age-matched controls.

**Figure 4.**
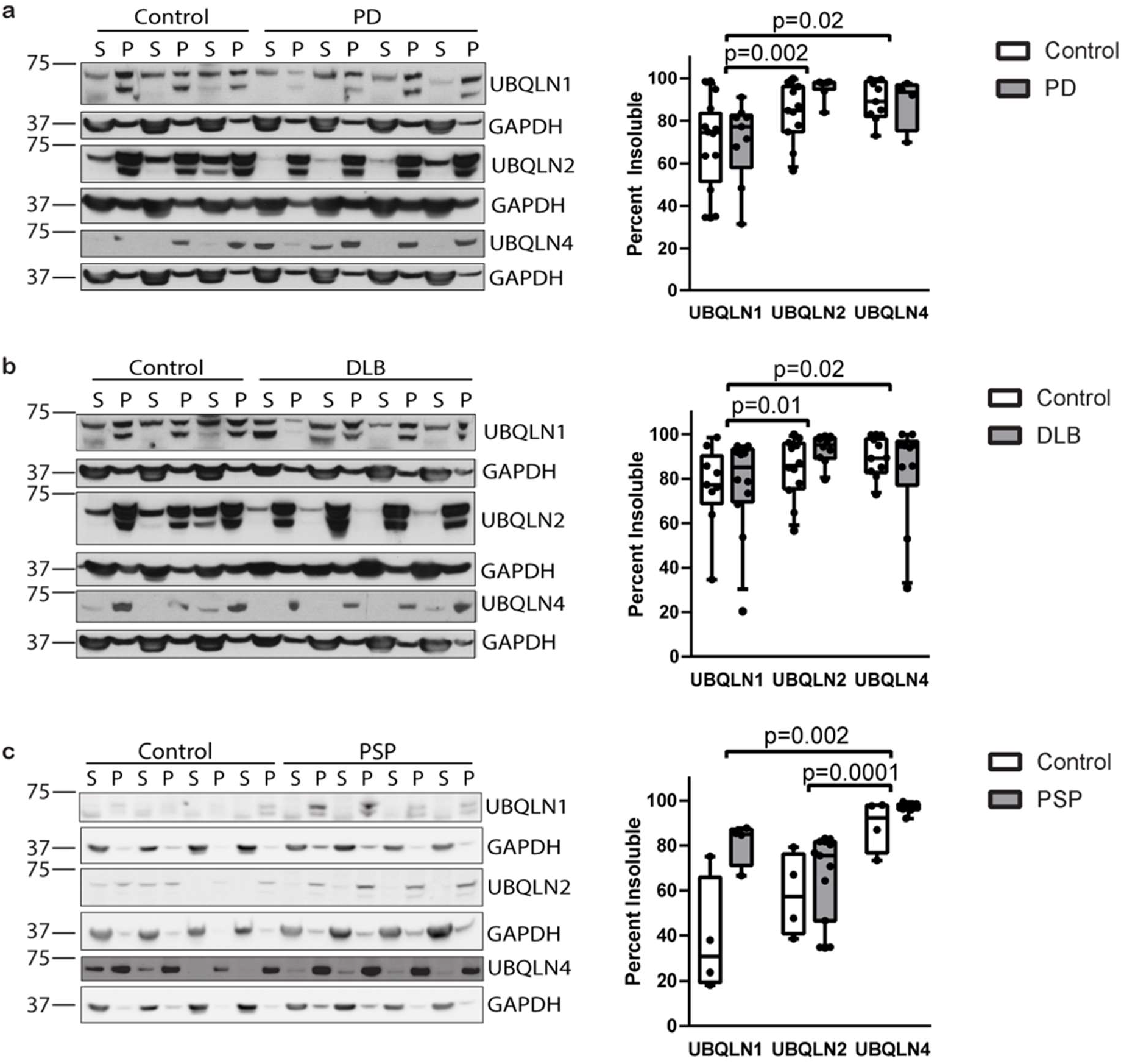
UBQLN2 and UBQLN4 are more insoluble than UBQLN1 in human brain tissue. Representative Western blots (uncropped Western blots shown in Supplemental Figure 3) of cingulate cortex from human controls (n=9), PD (**(a)**; n=6) and DLB (b) (; n=15), and mid-frontal gyrus from human controls (n=4) and PSP (**(c)**; n=12). Results display increased partitioning of UBQLN4 into the insoluble fraction compared to UBQLN1, analyzed by ANOVA grouping disease and controls together for each analysis. UBQLN2 insolubility is greater than UBQLN1 in both DLB and PD and respective controls, but significantly lower than UBQLN4 in PSP and respective controls. Results are expressed as a percentage of PBS-insoluble ubiquilin, determined from the calculated total PBS-insoluble (P) and PBS-soluble (S). No significant differences were seen between disease and controls for PD, DLB or PSP. Both UBQLN1 and UBQLN2 were detected and quantified as a doublet, as previously reported ^13,49,50^. Uncropped Western blots are shown in Supplemental Figure S3. Data were analyzed by one-way ANOVA using the Kruskal-Wallis (KW) post-hoc test. Box-and-whisker plot (median, first and third percentiles, range of quantified bands) is displayed with the scatter plot of the raw data.

No disease-dependent differences in UBQLN4 insolubility were detected in PD (Fig. 4a), DLB (Fig. 4b) or PSP (Fig. 4c) when compared to age-matched control samples. Thus, both in control and disease brain tissue, UBQLN4, and to a lesser extent UBQLN2, preferentially exists in an insoluble form.

### Divergence in ubiquilin mobility in condensates assessed by recovery after photobleaching

Using fluorescence recovery after photobleaching (FRAP), we previously showed that puncta of UBQLN2 in HEK-293T cells resemble liquid condensates and established a link between UBQLN2 condensate formation and aggregation ^13^. The differential aggregation propensity of the three ubiquilins prompted us to compare the mobility of each ubiquilin condensate in cells. To assess mobility, we used wild-type human ubiquilins fused to eGFP and measured FRAP of puncta in transfected cells. To reduce the chance that puncta exchanged contents with neighboring puncta, we examined their surroundings for other puncta using a Z-height scan. Data were recorded only from puncta that did not have immediately neighboring puncta and the bleached area was 80% or smaller than the total puncta area. We bleached only the center of each punctum and used fixed ROI across all puncta, so that fluorescence recovery that did occur would likely take place from nonbleached regions of the same punctum.

Consistent with the above results, the recovery time of eGFP-UBQLN1 fluorescence occurred more rapidly than that of UBQLN2 or UBQLN4 (Fig. 5c). To minimize the effect of puncta drift from the photobleached region (region of interest (ROI)), we chose shorter recording times of 300sec for UBQLN1 puncta, which were the fastest to recover signal after FRAP. Although the mobile fractions for all ubiquilins were similar, the mobile fractions for UBQLN2 and 4 trended lower than for UBQLN1 (Fig. 5d). Recovery times of UBQLN2 and 4 puncta are similar to one other and longer than those of UBQLN1 puncta (Fig. 5c). Taken together, these results suggest that UBQLN2 and UBQLN4 display similar self-association in liquid-like condensates, differing from UBQLN1.

**Figure 5.**
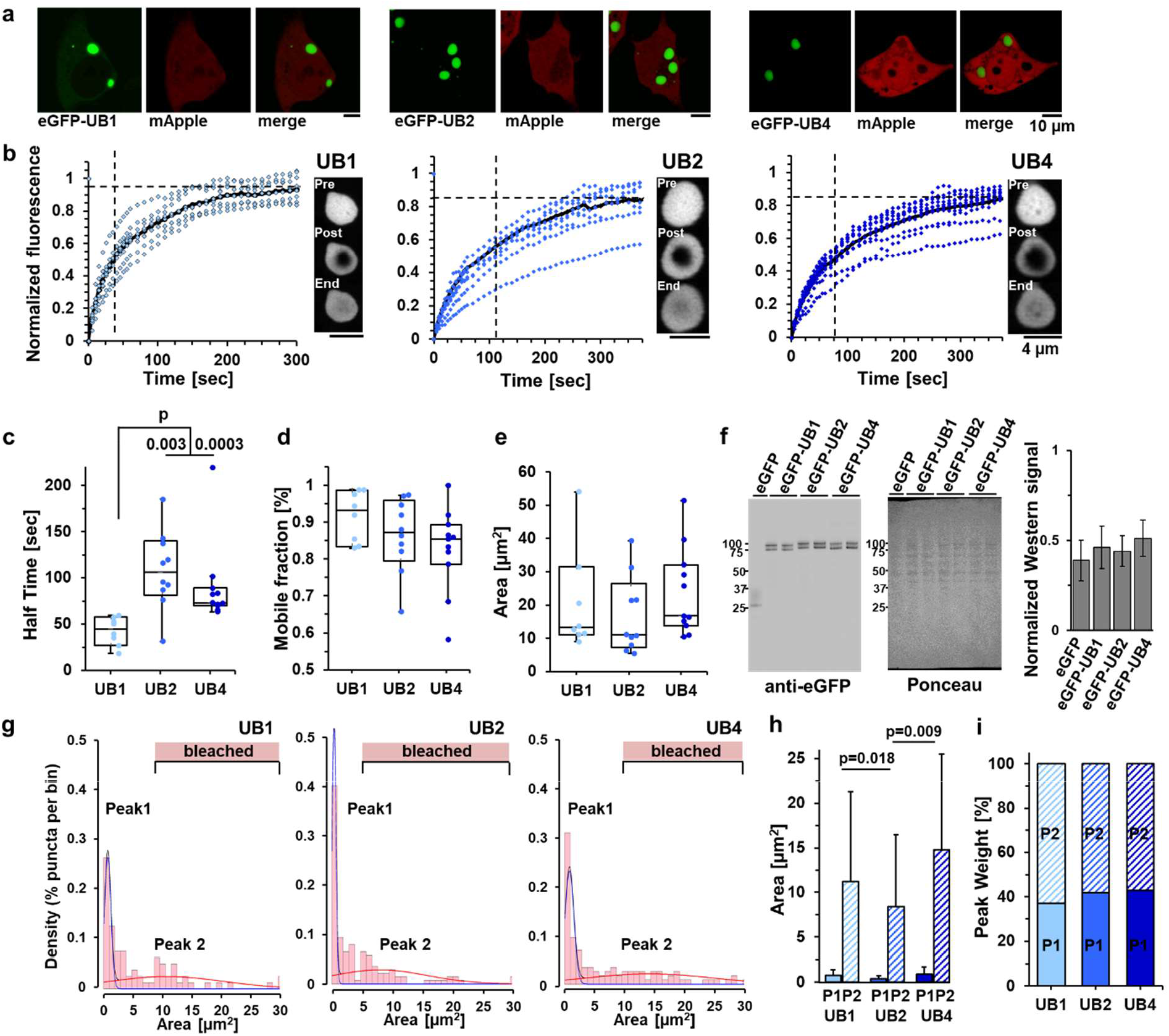
Dynamics of recovery after photobleaching of liquid-like ubiquilin condensates reveal divergent behavior of UBQLN2 and 4 versus UBQLN1. **(a)** Representative images of HEK-293T cells transfected with eGFP-UBQLNs 1, 2 or 4 and mApple. **(b)** The black curves represent the mean fluorescence recovery of eGFP-UBQLNs 1, 2 and 4 puncta over time. FRAP of the individual puncta is in blue. Dashed vertical and horizontal black lines are half-time and mobile-fraction means, respectively. Representative images of puncta immediately before (top) and after (middle) photobleaching, and at the end of data collection (bottom) are on the right of each plot. **(c)** Time to 50% fluorescence recovery after photobleaching is slower for eGFP-UBQLN2 and 4 than eGFP-UBQLN1. **(d)** FRAP measurements were used to compare protein mobility in the puncta. The brain-expressed ubiquilins have similar signal recovery (mobile fractions). Although not statistically significant, the mobile fractions of eGFP-UBQLN2 and 4, trended to be smaller than eGFP-UBQLN1. **(e)** The area of the puncta used for FRAP was similar for all three ubiquilins. **(f)** Representative Western blot (anti-eGFP antibody, left), and its Ponceau stain (total protein loaded, middle), and quantification (right) of lysed HEK293T transfected with eGFP-ubiquilins show that eGFP-tagged ubiquilins were expressed at similar levels (means, SEMa are displayed, N=4). **(g)** Area distribution of puncta was modeled with two Gaussians showing two populations of puncta (maximum mixed model likelihood: −251 for eGFP-UBQLN1, −377 for UBQLN2, and −375 for eGFP-UBQLN4). The pink bars labeled ‘bleached’ designate the size ranges of the bleached puncta. **(h)** Population P1 contains puncta of uniformly small area, and population P2 contains the remaining puncta spanning a wide range of large size areas. The punctum areas of eGFP-UBQLN1 and eGFP-UBQLN4 are larger than eGFP-UBQLN2. **(i)** Percent distribution (population weight) of the brain-expressed ubiquilins is 37-43% for P1, and 57-63% for P2. The data shown in **(g), (h)**, and **(i)** panels include puncta in the images taken from live HEK293-T cells transfected with eGFP-ubiquilins. Box-and-whisker plot (median, first and third percentiles, range of data) display the scatter plot of the raw data (N=5). Data were analyzed with the Kruskal-Wallis (KW) post-hoc test ^47^.

Modeling with normal (Gaussian) distributions revealed that the size of puncta formed by all three brain-expressed ubiquilins followed similar area distributions, consisting mainly of two populations (Fig. 5g-i). The first population, comprising 37-43% of puncta, was relatively small in size (mean areas: 0.8±0.6μm^2^ for eGFP-UBQLN1, 0.4±0.3 μm^2^ for eGFP-UBQLN2, and 0.9±0.8 μm^2^ for eGFP-UBQLN4), and the second population was much larger and distributed over a wider size range (mean areas: 11.2±10.1μm^2^ for eGFP-UBQLN1, 8.4±8.1μm^2^ for eGFP-UBQLN2, and 14.8±10.7 μm^2^ for eGFP-UBQLN4). Linear modeling of punctum area of live cells overexpressing ubiquilin proteins showed that larger puncta tend to be less circular (Supplemental Table S2, Fig. 5g-i). Overall, punctum areas of eGFP-UBQLN1 and eGFP-UBQLN4 were larger for both the small and large populations compared to eGFP-UBQLN2. We note that puncta selected for FRAP fall within the larger population of condensates (Fig. 5g-i).

## Discussion

All three brain-expressed ubiquilins are linked to neurodegenerative diseases characterized by protein dyshomeostasis ^1–3,5,28^. Recent studies highlight the propensity of UBQLN2 to form amyloid-like fibrils and liquid-like condensates ^13,19,20^, but the relative aggregation behavior of these three highly homologous proteins has not been investigated before. Here we report that the three ubiquilins share the property of liquid-liquid phase transition, yet differ in condensate properties (Fig. 5), propensity to form fibrillar aggregates (Fig. 2), and accumulation in insoluble species in cells and brain tissue (Figs. 3, and 4).

With respect to the shared property of liquid-liquid phase transition, UBQLN1 is most liquid-like and UBQLN4 most aggregation-prone among the three tested ubiquilins. Previous studies have suggested that liquid droplets formed by aggregation-prone proteins proceed to a less liquid state (from hydrogel to amyloid) at a faster rate ^33^. Consistent with our earlier findings with UBQLN2 and studies by others of additional aggregation-prone proteins ^13,15,17,34^, the increased recovery times after photobleaching for UBQLN4 puncta (Fig. 5) correlate with its tendency to form high molecular weight aggregates and partition into the insoluble fraction (Figs. 3, and 4). Some studies have shown that amyloid fibrils can induce liquid-liquid phase transition, implying that the relationship could be bi-directional ^35,36^. The correlation between increased recovery times of UBQLN4 liquid droplets and heightened fibril formation suggests that regulation of liquid-liquid phase transition by ubiquilins is linked to aggregation.

The ability of all ubiquilins to form liquid droplets suggests it is a fundamental shared principle of this class of proteins that relates, in a still undefined manner, to their function in quality control pathways. While the relevance of liquid droplet formation to ubiquilin function is not yet understood, studies of other protein quality control factors may provide insight into potential mechanisms. For example, proteasome components form liquid droplets that depend on the interaction of polyubiquitin chains with the UBA-UBL ubiquitin-proteasome shuttle factor, Rad23B (38). Indeed, Alexander et al ^25^ showed that UBQLN2 is involved in the regulation of stress granules. Thus, ubiquilins form concentrated molecular assemblies possibly *via* their UBL and UBA domains to regulate multiple cellular processes.

Despite being highly similar proteins, the brain-expressed ubiquilins differ in condensate and aggregation behavior. These differences suggest that relatively minor sequence changes can alter the behavior of the protein. In this light, one can envision that missense mutations that cause UBQLN2-mediated disease, or possibly UBQLN4-mediated disease, would change behavior of these proteins along the liquid droplet-hydrogel-aggregate spectrum. Published evidence from our group and others supports this view ^13,19,20^. While the UBL and UBA domains of ubiquilins are nearly identical, the three brain-expressed UBQLNs are distinguished by the middle region located between the UBL and UBA domains ^29^. In evaluating the C-terminal regions of UBQLNs 1, 2 and 4 for fibril formation, we demonstrated that UBQLN4^444-601^ was the most aggregation-prone, while UBQLN1^438-589^ and UBQLN2^430-624^ displayed slower kinetics of aggregation. These results suggest that C-terminal differences among these three brain-expressed ubiquilins lead to altered aggregation behavior that corresponds to the differences observed for the full-length proteins.

Although UBQLN4 exhibited the greatest propensity to aggregate in *in vitro* assays, cellular models, and human brain tissue, UBQLN2 also displayed heightened aggregation propensity in cellular models and human brain when compared to UBQLN1. Since UBQLN2 is known to accumulate in aggregate-like structures in various neurodegenerative diseases ^3,28,37^, it will be important now to determine whether UBQLN4 similarly accumulates in one or more neurodegenerative diseases.

Our results demonstrating that UBQLN4 insolubility is elevated over UBQLN1—and in some cases over UBQLN2—in human brain suggest that heightened UBQLN4 aggregation behavior may be an intrinsic feature of the protein that informs its function. Furthermore, the percent UBQLN4 in the insoluble fraction showed little variability between samples in spite of the inherent wide variation present when using human tissue. While no differences in UBQLN4 solubility were detected in PD, DLB or PSP samples compared to age-matched controls, our analysis of available human samples is limited and does not allow us to conclude that there are no disease-dependent changes. Further studies are needed to elucidate whether the aggregation status of UBQLN4 relates to specific diseases or more generally to brain aging.

Although the relationship between formation of dynamic molecular assemblies and the function of ubiquilins is yet to be clarified, we anticipate that the phase separation and aggregation behavior of all three brain-expressed ubiquilins will prove to be regulated by ubiquitin and will inform their function as proteostasis regulatory factors ^19^. Our results reveal two populations of condensates formed by all three ubiquilins, one being uniformly small and the second being larger and more variable in size. It is possible that small condensates represent functional assemblies, whereas larger condensates do not. Studies have also shown that the three ubiquilins interact with one another, indicating that their cellular functions and liquid-liquid phase transition properties may be linked ^38^. Further research will be needed to better understand how the shared and divergent condensate and aggregation properties of ubiquilins influence their cellular functions and roles in disease.

## Methods

### Plasmids

The pCMV4-FLAG-UBQLN2 plasmid (p4455 FLAG-hPLIC-2; Addgene plasmid # 8661) and pCS2-FLAG-UBQLN1 plasmid (p4458 FLAG-hPLIC-1; Addgene plasmid # 8663) were gifts from Peter Howley (42). UBQLN4 was cloned from pDONR223-UBQLN4 (pENTR-A1UP; Addgene plasmid # 16170), which was a gift from Huda Zoghbi (Baylor College of Medicine), into the pCMV4-FLAG vector. Control empty vector plasmid for FLAG cell transfection experiments was pCMV-HA. eGFP-UBQLN1, 2 and 4 were cloned from FLAG-tagged plasmids and eGFP. The pRP[Exp]-Neo-CMV-eGFP, pRP[Exp]-Neo-CMV-eGFP-UBQLN1, pRP[Exp]-Neo-CMV-eGFP-UBQLN2, and pRP[Exp]-Neo-CMV-eGFP-UBQLN4 were purchased from VectorBuilder. The plasmids for bacterial expression of ubiquilin constructs (pET3a-His6-W-UBQLN1^438-589^, pET3a-His6-W-UBQLN2^430-624^, pET3a-His6-W-UBQLN4^444-601^ pET3a-His6-W-UBQLN2^581-624^(UBA)) were purchased from Genscript.

### Protein expression and purification

All plasmids for bacterial expression contained a 6xHis and a tryptophan (Trp) at the N-terminus. UBQLN1^438-589^, UBQLN2^430-624^ and UBQLN4^444-601^ constructs were transformed in Rosetta (DE3) Escherichia coli bacteria. All LB agar plates and LB media were supplemented with 100 μg/ml carbenicillin and 34 μg/ml chloramphenicol. Transformed cells were grown overnight at 37°C on LB agar media plates. The following day, cells were transferred to 100 mL LB starter cultures and allowed to grow for 60 min, then upscaled into 1L LB media. At OD601 ≈ 0.6-0.8, cells were induced with 0.5 mM isopropyl β-D-1-thio-galacto-pyranoside (IPTG) and collected after 4 additional hours of incubation. Bacteria were collected by centrifugation for 10 min at 10,322 x g. Bacterial pellets were stored at −80°C.

For protein purification, pellets of bacteria grown in 1 L of LB media were resuspended in 25 ml of pre-chilled lysis buffer containing 2% glycerol, 1 mM EDTA, 25 mM Na phosphate, pH 7.4, EDTA-free cOmplete protease inhibitor cocktail tablet (Roche; 1 tablet per 10mL lysis buffer) and 6 μL/mL of saturated phenylmethylsulfonyl fluoride (PMSF). Bacteria were then lysed using EmulsiFlex B-15 high pressure homogenizer (Avestin). The lysates were centrifuged at 31,000 x g for 20 min and the supernatant was added to Ni-NTA agarose slurry, After incubation for 15 min at 4° C, the slurry was washed with 25 mM Na phosphate pH 7.4 containing 0.1M NaCl, 6 ul/ml from saturated PMSF 2% glycerol, and 20mM imidazole. Proteins were eluted with 25mM Na phosphate pH 7.4, which was supplemented with 200 mM imidazole 0.1 M NaCl, 6 μl/ml from saturated PMSF, and 2% glycerol. In the case UBQLN2430-624 construct, the eluted protein was diluted with two volumes of 25 mM Na phosphate pH 7.4 containing 2% glycerol. Then it was filtered through 50kDa concentrator (Amicon Ultra-15 Centrifugal Filter Unit, Millipore Sigma) and the flow through was collected for dialysis. Proteins were dialyzed against 5 mM Na phosphate pH 7.4 with two buffer exchanges overnight. Immediately after dialysis, proteins were frozen in liquid nitrogen and stored at −80°C. Protein concentration was determined by Pierce BCA protein assay Kit (cat# 23225, ThermoFisher Scientific).

### ThioflavinT binding assay

ThT assays were carried out with 10 μM protein in 0.1 M NaCl, 1 mM sodium azide, and 20 mM Na phosphate pH 7.5. Prior to the assay, proteins were filtered through a 0.22 μm filter. ThT was added to a final concentration of 10 μM. Teflon beads were added into each well of a Falcon 96-well plate (black/clear, flat bottom, Corning, cat # 353219). Then 75 μl of sample were pipetted into each well. Plates were then incubated at 37°C in a FLUOstar Omega (BMG Labtech Inc) by shaking at 200 rpm using the ‘meander corner well shaking’ mode. Fluorescence was measured with gain set at 90%, an excitation wavelength of 440 nm and emission wavelength of 490 nm. Three technical replicates were measured per sample for a single ThT assay. Data shown in Fig. 1 are averaged over three to four independent experiments done with different protein preparations. Half times (50% steady-state transition) were calculated as described by ^39^.

### Transmission electron microscopy (TEM)

Negatively stained specimens for TEM were prepared by applying 5 μL of protein sample to hydrophilic 400 mesh carbon-coated Formvar support films mounted on copper grids (Ted Pella, Inc., 01702-F). Samples were allowed to adhere for 4 min, rinsed twice with distilled water, and stained for 60-90 sec with 5 μL of 1% uranyl acetate (Ted Pella, Inc.). All samples were imaged at an accelerating voltage of 80 kV in a JEOL JSM 1400 Plus (JOEL).

### HEK cell transfection

Human embryonic kidney 293 (HEK-293; Batch #70008735, ATCC) cells were cultured in high glucose DMEM, supplemented with 10% FBS, 10mM Glutamine and 100 U/ml penicillin/streptomycin. Cells were transfected with either pCMV-FLAG-UBQLN1, pCMV4-FLAG-UBQLN2, pCMV4-FLAG-UBQLN4 or pCMV-HA using Lipofectamine-2000 according to the manufacturer’s instructions. HEK-293T (cat# CRL 3216, ATCC Batch No.70008735) cultured in DMEM (cat# SH30242.01 ThermoFisher Scientific), 10% FBS and 100 U/ml penicillin/streptomycin, were co-transfected as with constructs of eGFP fusion proteins (proteins as specified), and mApple (for visualization of cells).

### Human disease brain tissue

Frozen brain tissue from the cingulate cortex was obtained from subjects with PD, DLB, and age-matched control subjects as well as mid-frontal cortex from PSP and age-matched control subjects (Supplemental Table S1) from the Michigan Brain Bank (University of Michigan, Ann Arbor, MI, USA). Brain tissue was collected with the informed consent of the patients. Protocols were approved by the Institutional Review Board of the University of Michigan and abide by the Declaration of Helsinki principles. Samples were examined at autopsy by neuropathologists for diagnosis.

### Mouse Models

This study was conducted in a facility approved by the American Association for the Accreditation of Laboratory Animal Care, and all experiments were performed in accordance with the National Institutes of Health Guide for the Care and Use of Laboratory Animals and approved by the Institutional Animal Care and Use Committee of the University of Michigan. C57/Bl6 mice were housed at the University of Michigan animal care facility and maintained according to U.S. Department of Agriculture standards (12 h light/dark cycle with food and water available ad libitum).

### Western blot analysis

Cell lysates were homogenized using 0.2 mm stainless steel beads, sonicated for 5 minutes in chilled water, centrifuged at 10000 rcf for 10 min at 4°C and supernatants were collected. For insoluble fractions, pellets were resuspended in PBS with protease inhibitor cocktail (catalog no. 11873580001; Sigma Aldrich), centrifuged at 10000 rcf for 10 min at 4°C, and supernatants were discarded. Remaining pellet was resuspended in 1% sarkosyl in PBS with protease inhibitor, vortexed for 1 min, and incubated at room temperature for 30 minutes. Samples were water sonicated for 5 min and centrifuged for 20 min at 14000 rpm at 4°C. Protein concentrations were measured by BCA (cat#23227, ThermoScientific). Cell lysates containing 10 μg of total protein were loaded (without boiling) on precast NuPAGE 4-12% Bis-Tris gels (Invitrogen) for SDS-PAGE analysis. Gels were subsequently transferred onto nitrocellulose membranes and stained with Ponceau-S for total protein quantification. After destaining, membranes were blocked for 1 hour at room temperature with 10% nonfat dry milk in TBS-T buffer. Membranes were then probed overnight at 4℃ in anti-FLAG, clone M2 (Sigma, cat. #F3165; 1:1000), anti-Ubiquilin-2 (Novus Biologicals, cat. #NBP2-25164; 1:2000), anti-Ubiquilin-1 (Novus biologicals, cat. #H00029979-M02; 1:2000) or anti-A1UP (Santa Cruz, cat. #sc-136145; 1:2000) diluted in 5% nonfat dry milk. HRP-conjugated goat anti-rabbit IgG or goat anti-mouse IgG (1:5000; ThermoFisher Scientific, cat. # 31460 and 32430) were used for detection as appropriate. All ubiquilin antibodies were tested without stripping. In replicate Western blots, secondary antibodies were inactivated by incubating with H2O2 for 15 minutes at 37C before adding another antibody of a different species. ECL (Pierce) was used to visualize bands using the Gbox Mini-6 (Syngene), which were normalized to corresponding total protein levels detected by Ponceau-S. All quantification of immunoblots was performed by densitometric analysis using Genetools software (Syngene). Analyses were completed in triplicate and analyzed by one-way ANOVA with the Bonferroni post-hoc test.

HEK-293T cells (cat# CRL 3216, ATCC Batch No.70008735) transfected with eGFP tagged ubiquilins were resuspended in ice cold PBS 0.5% Triton-X 100 with cOmplete™, Mini, EDTA-free Protease Inhibitor Cocktail (cat# 11836170001, Millipore Sigma/Roche), and sonicated in Bioruptor Pico (Diagenode) using 4 sonication cycles (each 30 sec ON/30 sec OFF) at 4°C. After centrifugation at 10000 rcf for 10 min at 4°C, supernatants were snap frozen in liquid nitrogen and stored at −80°C until use. Pierce BCA protein assay Kit (cat# 23225, ThermoFisher Scientific). Samples containing 5.5 μg of total protein supplemented with NuPage LDS sample buffer (cat# NP0007, ThermoFisher Scientific) were loaded without boiling on NuPAGE 4-12% Bis-Tris gels (cat# WG1403BOX, ThermoFisher Scientific) with running 1x NuPage MES SDS Running Buffer (cat# NP0002, ThermoFisher Scientific)for SDS-PAGE analysis. Gels were subsequently transferred onto nitrocellulose membranes (0.45 μm, cat# 162-0115, BioRad) for 1 hour at 110V and stained with Ponceau-S for total protein quantification. After destaining, membranes were blocked for 1 hour at room temperature with 10% Applichem nonfat dried milk (cat# NC0167677, ThermoFisher Scientific) in TBS-T buffer. Membranes were then probed overnight at 4℃ in anti-GFP (mouse monoclonal IgG1κ, cat#: 11814460001 Millipore Sigma/Roche) diluted 1:1000 in 5% Applichem nonfat dried milk in TBS-T. HRP-conjugated goat anti-mouse IgG (diluted 1:3000 in 5% Applichem nonfat dried milk in TBS-T) were used for detection. Advansta WesternBright ECL HRP Substrate Kits (cat# 490005-020, WVR) were used to visualize protein bands. Protein bands of five independent experiments were quantified using ImageJ ^40^ and normalized for the total protein amount detected by Ponceau-S.

Soluble and insoluble human brain samples and mouse brain samples were homogenized in PBS with a protease inhibitor cocktail (catalog no. 11873580001; Sigma Aldrich) with 3.2 mm stainless steel beads, using a 1:3 dilution of tissue: PBS (w/v). Samples were centrifuged at 10000 rcf for 10 min at 4°C. Supernatants (PBS-soluble fraction) were aliquoted, snap-frozen, and stored at −80°C until use. For insoluble fractions, pellets were resuspended in PBS with the protease inhibitor cocktail, centrifuged at 10000 rcf for 10 min at 4°C and supernatants were discarded. The remaining pellet was resuspended in 1% sarkosyl in PBS with protease inhibitor, vortexed for 1 min, and incubated at room temperature for 1 hr. Samples were water sonicated for 5 min and centrifuged for 20 min at 14000 rpm at 4°C. Supernatants were discarded and procedure was repeated with remaining pellet for insoluble fraction. Protein concentrations were measured by BCA (cat#23227, ThermoScientific). Brain extracts containing 25 μg of total protein were loaded (without boiling) on precast NuPAGE 4-12% Bis-Tris gels (Invitrogen) for SDS-PAGE analysis. Gels were subsequently transferred onto nitrocellulose membranes and blocked for 1 hour at room temperature with 10% nonfat dry milk in TBS-T buffer. Membranes were then probed overnight at 4℃ in anti-Ubiquilin-2 (Novus Biologicals, cat. #NBP2-25164; 1:2000), anti-Ubiquilin-1 (Novus biologicals, cat. #H00029979-M02; 1:2000), anti-A1UP for UBQLN4 (Santa Cruz, cat. #sc-136145; 1:2000), anti-Histone H3 (Cell Signaling, cat. #4499; 1:2000), anti-β-actin (Sigma, cat. #AC15; 1:5000) or anti-GAPDH (Millipore, cat. #MAB374; 1:5000) diluted in 5% nonfat dry milk. HRP-conjugated goat anti-rabbit IgG or goat anti-mouse IgG (1:5000; ThermoFisher Scientific, cat. # 31460 and 32430) were used for detection as appropriate. ECL (Pierce) was used to visualize bands using the Gbox Mini-6 (Syngene), which were normalized to corresponding GAPDH levels. All quantification of immunoblots was performed by densitometric analysis using ImageJ software (National Institutes of Health) ^40^.

To quantify the percent insoluble protein for each sample, normalized insoluble ubiquilin band densities were corrected for ubiquilin concentration based on lysate volume, and then divided by total (soluble + insoluble) normalized ubiquilin protein levels and multiplied by 100. Two-way ANOVA revealed UBQLN-dependent differences, but no differences based on disease state. Therefore, control and disease samples were pooled for one-way ANOVA analyses of differences in UBQLN insolubility. Analyses were completed in triplicate and analyzed by one-way ANOVA with the Kruskal-Wallis post-hoc test.

### Immunofluorescence

Fixed cells were washed in PBS, permeabilized with 0.5% Triton-X 100 and blocked in 5% goat serum for 1 hour. Cells were incubated in anti-FLAG, clone M2 (Sigma cat. #F3165, 1:100) overnight at 4°C. The following day, cells were washed in PBS three times for 10 min each and incubated with goat anti-mouse IgG Alexa-568 (Invitrogen cat. #A-11004; 1:500) for 1 h. Sections were then washed in PBS three times for 10 min each and incubated with DAPI (Sigma) to label nuclei for 5 min at room temperature, washed three times for 5 min each, and were mounted with Prolong Gold Antifade Reagent (Invitrogen). Slides were imaged using an IX71 Olympus inverted microscope. Images were analyzed with the Analyze Particles tool in Image-J (National Institute of Health) (39) to determine the corrected total cellular fluorescence (CTCF; used as a measure of overall expression of each ubiquilin protein per cell for normalization) and the corrected total fluorescence for each punctum (CTFP). To calculate the percent fluorescence of puncta, the sum of CTFP of all puncta in the cell was divided by CTCF of the whole cell. Immunofluorescence experiments were repeated in triplicate and statistical analyses utilized means from the three replicates. Analyses were completed by one-way ANOVA and the Bonferroni post-hoc test.

### Fluorescence recovery after photobleaching (FRAP)

HEK293-T cells (Batch #70008735, ATCC) were plated on Nunc™ Lab-Tek™ II Chambered Coverglass (cat. #155409, Fisher) in DMEM, supplemented with 10% FBS, 10 mM Glutamine and 100 U/ml penicillin/streptomycin. The next day cells were transfected with eGFP-UBQLN1, eGFP-UBQLN2, and eGFP-UBQLN4 plasmid DNA. Using Lipofectamine™ 2000 Transfection Reagent (cat# 11668027; Fisher) according to the manufacturer’s instructions. Cells were imaged 24 hours after transfection with a Nikon A-1 confocal microscope (40x WI Lens) using Nikon Elements software with perfect focus engaged. For FRAP data collection, an area of 512 pixels was scanned using speed 518 frames/sec. FRAP imaging consisted of three phases: pre-bleach, bleaching, and post-bleach imaging. During pre-bleach imaging, each punctum was imaged every 2 seconds for 10 seconds. In the bleach phase, a 5-6 μm^2^ ROI was drawn in the middle of the punctum covering no more than 2/3. This ROI defined the stimulation area for a 488 nm laser with 20% power. The post-bleach phase consisted of two periods. For the first minute, images were acquired every 5 seconds, while for the subsequent 6 minutes, images were acquired every 10 seconds. This was done in order to provide greater temporal resolution during the rising phase of fluorescence recovery. FRAP analysis was performed in Fiji ^41^. To fix granules in place, stack registration (Rigid Body) was performed. Following thresholding, one ROI was generated that corresponded to the pre-bleach signal and another to the postbleach signal. A mask was created from the pre-bleach ROI, inverted, and the postbleach ROI was subtracted from this image. This created a region of bleached signal deemed the FRAP ROI. At each time point the integrated density of the FRAP ROI was divided by the integrated density of the pre-bleach ROI. These values were then normalized such that the mean of the five pre-bleach values was set to 1, and the first postbleach value was set to 0. To calculate the mobile fraction and half time, the normalized fluorescent signal measured from each granule was fitted to the equation A*(1-exp(-t/τ)), where A is the mobile fraction, t is the collection time, τ is the half time ^42^. Puncta area, circularity, roundedness, aspect ratio. and solidity were calculated using Fiji ^41^.

The area distribution of UBQLN1, 2, and 4 puncta was modeled using Normal (μ, σ2) distribution. The process of distribution model-fitting involved maximum likelihood estimation of the model distribution parameters ^43^. We employed the R packages mixtools ^44^ and ggplot2 ^45^ to obtain and plot the desired models. The model parameters and the mixture distribution weights are used as proxy measures to discriminate between different puncta areas. For statistical analysis of FRAP, Kruskal-Wallis (KW) test was used, which is an alternative to One-Way Analysis of Variance (ANOVA) when parametric assumptions are not guaranteed ^46,47^. FRAP data are shown as means +/− SEM.

### Experimental design and statistical analysis

The accepted level of significance for all analyses was p ≤ 0.05. Data are expressed as means +/− SEM. P-values for overall ANOVAs are displayed in analyses that did not show a significant difference and individual post-hoc comparison P-values are displayed for significant ANOVAs. Data were analyzed using R (cran.r-project.org) and Statview (SAS Institute). Specific statistical analysis descriptions are detailed within each individual experimental method

## Contributions

JEG, HLP, and MII conceptualized and initiated the project; JEG, HLP, and MII designed experiments; JEG, HL, JX, ABS, FSK, NS, JR, NL, and MII performed experiments and acquired data, JEG, HL, ABS, LMS, NS, JR, NL, and MII analyzed and interpreted data, JEG, FSK, HLP and MII wrote the manuscript; all authors critically read and edited the manuscript

## Acknowledgments

We thank Peter Howley and Huda Zoghbi for providing constructs, and Paulson lab members for their helpful suggestions. We thank Matthew Perkins for providing the human brain tissue from the UM Brain Bank, affiliated with the Michigan Alzheimer’s Disease Research Center.

## References

1 Deng, H.-X. et al. Mutations in UBQLN2 cause dominant X-linked juvenile and adult-onset ALS and ALS/dementia. Nature 477, 211–215 (2011).

2 Fahed, A. C. et al. UBQLN2 mutation causing heterogeneous X-linked dominant neurodegeneration. Annals of neurology 75, 793–798 (2014).

3 Mori, F. et al. Ubiquilin immunoreactivity in cytoplasmic and nuclear inclusions in synucleinopathies, polyglutamine diseases and intranuclear inclusion body disease. Acta neuropathologica 124, 149–151 (2012).

4 Rothenberg, C. & Monteiro, M. J. Ubiquilin at a crossroads in protein degradation pathways. Autophagy 6, 979–980 (2010).

5 Edens, B. M. et al. A novel ALS-associated variant in UBQLN4 regulates motor axon morphogenesis. Elife 6, e25453 (2017).

6 Elsasser, S. et al. Proteasome subunit Rpn1 binds ubiquitin-like protein domains. Nature cell biology 4, 725–730 (2002).

7 Rao, H. & Sastry, A. Recognition of specific ubiquitin conjugates is important for the proteolytic functions of the ubiquitin-associated domain proteins Dsk2 and Rad23. Journal of Biological Chemistry 277, 11691–11695 (2002).

8 Lim, P. J. et al. Ubiquilin and p97/VCP bind erasin, forming a complex involved in ERAD. Journal of Cell Biology 187, 201–217 (2009).

9 Chen, T., Huang, B., Shi, X., Gao, L. & Huang, C. Mutant UBQLN2 P497H in motor neurons leads to ALS-like phenotypes and defective autophagy in rats. Acta neuropathologica communications 6, 1–15 (2018).

10 Rothenberg, C. et al. Ubiquilin functions in autophagy and is degraded by chaperone-mediated autophagy. Human molecular genetics 19, 3219–3232 (2010).

11 Şentürk, M. et al. Ubiquilins regulate autophagic flux through mTOR signalling and lysosomal acidification. Nature cell biology 21, 384–396 (2019).

12 Lee, D. Y., Arnott, D. & Brown, E. J. Ubiquilin4 is an adaptor protein that recruits Ubiquilin1 to the autophagy machinery. EMBO Rep 14, 373–381, doi:10.1038/embor.2013.22 (2013).

13 Sharkey, L. M. et al. Mutant UBQLN2 promotes toxicity by modulating intrinsic self-assembly. Proceedings of the National Academy of Sciences 115, E10495–E10504 (2018).

14 Wegmann, S. et al. Tau protein liquid–liquid phase separation can initiate tau aggregation. The EMBO journal 37 (2018).

15 Patel, A. et al. A liquid-to-solid phase transition of the ALS protein FUS accelerated by disease mutation. Cell 162, 1066–1077 (2015).

16 Molliex, A. et al. Phase separation by low complexity domains promotes stress granule assembly and drives pathological fibrillization. Cell 163, 123–133 (2015).

17 Conicella, A. E., Zerze, G. H., Mittal, J. & Fawzi, N. L. ALS mutations disrupt phase separation mediated by α-helical structure in the TDP-43 low-complexity C-terminal domain. Structure 24, 1537–1549 (2016).

18 Mackenzie, I. R. et al. TIA1 mutations in amyotrophic lateral sclerosis and frontotemporal dementia promote phase separation and alter stress granule dynamics. Neuron 95, 808–816. e809 (2017).

19 Dao, T. P. et al. Ubiquitin modulates liquid-liquid phase separation of UBQLN2 via disruption of multivalent interactions. Molecular cell 69, 965–978. e966 (2018).

20 Dao, T. P. et al. ALS-linked mutations affect UBQLN2 oligomerization and phase separation in a position-and amino acid-dependent manner. Structure 27, 937–951. e935 (2019).

21 Fujioka, Y. et al. Phase separation organizes the site of autophagosome formation. Nature, 1–5 (2020).

22 Hyman, A. A., Weber, C. A. & Jülicher, F. Liquid-liquid phase separation in biology. Annual review of cell and developmental biology 30, 39–58 (2014).

23 Strom, A. R. & Brangwynne, C. P. The liquid nucleome–phase transitions in the nucleus at a glance. Journal of cell science 132 (2019).

24 Rhine, K., Vidaurre, V. & Myong, S. RNA Droplets. Annual Review of Biophysics 49 (2020).

25 Alexander, E. J. et al. Ubiquilin 2 modulates ALS/FTD-linked FUS–RNA complex dynamics and stress granule formation. Proceedings of the National Academy of Sciences 115, E11485–E11494 (2018).

26 Elbaum-Garfinkle, S. Matter over mind: Liquid phase separation and neurodegeneration. Journal of Biological Chemistry 294, 7160–7168 (2019).

27 Babinchak, W. M. & Surewicz, W. K. Liquid–Liquid Phase Separation and its Mechanistic Role in Pathological Protein Aggregation. Journal of Molecular Biology (2020).

28 Rutherford, N. J. et al. Unbiased screen reveals ubiquilin-1 and-2 highly associated with huntingtin inclusions. Brain research 1524, 62–73 (2013).

29 Marín, I. The ubiquilin gene family: evolutionary patterns and functional insights. BMC evolutionary biology 14, 63 (2014).

30 Goldschmidt, L., Teng, P. K., Riek, R. & Eisenberg, D. Identifying the amylome, proteins capable of forming amyloid-like fibrils. Proceedings of the National Academy of Sciences 107, 3487–3492 (2010).

31 Biancalana, M. & Koide, S. Molecular mechanism of Thioflavin-T binding to amyloid fibrils. Biochimica et Biophysica Acta (BBA)-Proteins and Proteomics 1804, 1405–1412 (2010).

32 Groenning, M. Binding mode of Thioflavin T and other molecular probes in the context of amyloid fibrils—current status. Journal of chemical biology 3, 1–18 (2010).

33 Lin, Y., Protter, D. S., Rosen, M. K. & Parker, R. Formation and maturation of phase-separated liquid droplets by RNA-binding proteins. Molecular cell 60, 208–219 (2015).

34 Murakami, T. et al. ALS/FTD mutation-induced phase transition of FUS liquid droplets and reversible hydrogels into irreversible hydrogels impairs RNP granule function. Neuron 88, 678–690 (2015).

35 Xiang, S. et al. The LC domain of hnRNPA2 adopts similar conformations in hydrogel polymers, liquid-like droplets, and nuclei. Cell 163, 829–839 (2015).

36 Gui, X. et al. Structural basis for reversible amyloids of hnRNPA1 elucidates their role in stress granule assembly. Nature communications 10, 1–12 (2019).

37 Zeng, L. et al. Differential recruitment of UBQLN2 to nuclear inclusions in the polyglutamine diseases HD and SCA3. Neurobiology of disease 82, 281–288 (2015).

38 Hjerpe, R. et al. UBQLN2 mediates autophagy-independent protein aggregate clearance by the proteasome. Cell 166, 935–949 (2016).

39 Arosio, P., Knowles, T. P. & Linse, S. On the lag phase in amyloid fibril formation. Physical Chemistry Chemical Physics 17, 7606–7618 (2015).

40 Rueden, C. T. et al. ImageJ2: ImageJ for the next generation of scientific image data. BMC bioinformatics 18, 529 (2017).

41 Schindelin, J.

42 Axelrod, D., Koppel, D., Schlessinger, J., Elson, E. & Webb, W. W. Mobility measurement by analysis of fluorescence photobleaching recovery kinetics. Biophysical journal 16, 1055 (1976).

43 Dinov, I. D. Expectation maximization and mixture modeling tutorial. (2008).

44 Benaglia, T., Chauveau, D., Hunter, D. & Young, D. mixtools: An R package for analyzing finite mixture models. (2009).

45 Wickham, H. ggplot2: elegant graphics for data analysis. (Springer, 2016).

46 Conover, W. J. & Conover, W. J. Practical nonparametric statistics. (1980).

47 Chu, A., Cui, J. & Dinov, I. D. SOCR analyses: implementation and demonstration of a new graphical statistics educational toolkit. Journal of statistical software 30, 1 (2009).

48 Robert, X. & Gouet, P. Deciphering key features in protein structures with the new ENDscript server. Nucleic acids research 42, W320–W324 (2014).

49 Mah, A. L., Perry, G., Smith, M. A. & Monteiro, M. J. Identification of ubiquilin, a novel presenilin interactor that increases presenilin protein accumulation. The Journal of cell biology 151, 847–862 (2000).

50 Kim, S. H. et al. Mutation-dependent aggregation and toxicity in a Drosophila model for UBQLN2-associated ALS. Human molecular genetics 27, 322–337 (2018).

