## Supporting Information for "Shared and divergent phase separation and aggregation properties of brain-expressed ubiquilins"

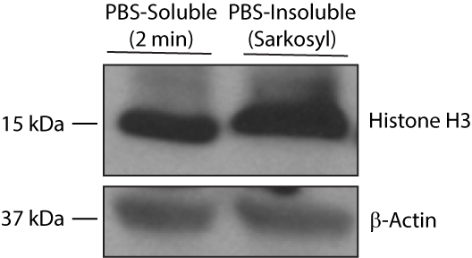

**Supplementary Figure S1.**

Homogenization of 3-month-old C57/Bl-6 mouse brain for only 2 minutes (less than 20% of the time we use in our protocols to homogenize tissue) in a detergent-free buffer (1x PBS) is sufficient to release nuclear (Histone H3) and cytoplasmic (β-actin) protein into the PBS-soluble fraction when run alongside the PBS-insoluble pellet lysed in a denaturing buffer (Sarkosyl).

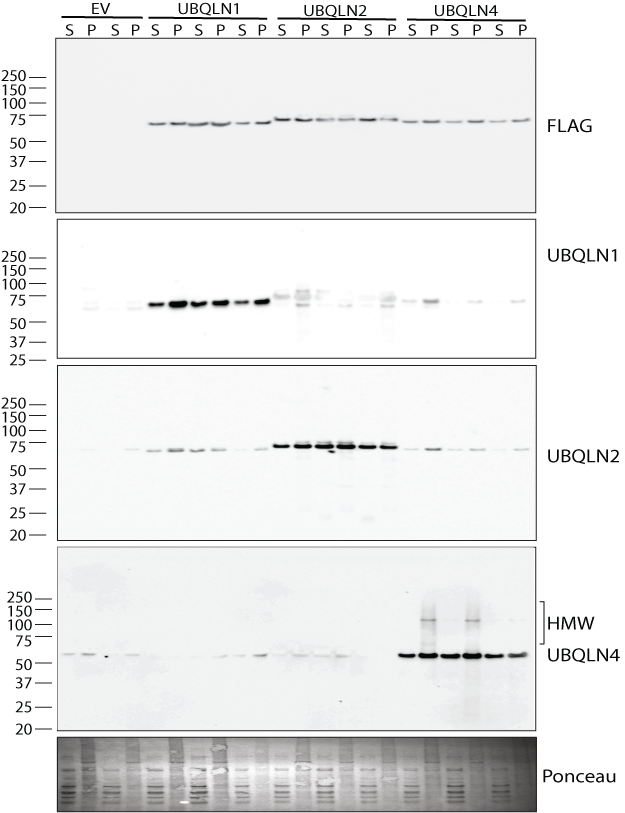

**Supplementary Figure S2.**

Full, uncropped Western blots of cell lysates from Figure 3 detected with UBQLN1, 2 and 4-specific antibodies demonstrate specificity of each antibody in measuring the ubiquilin proteins.

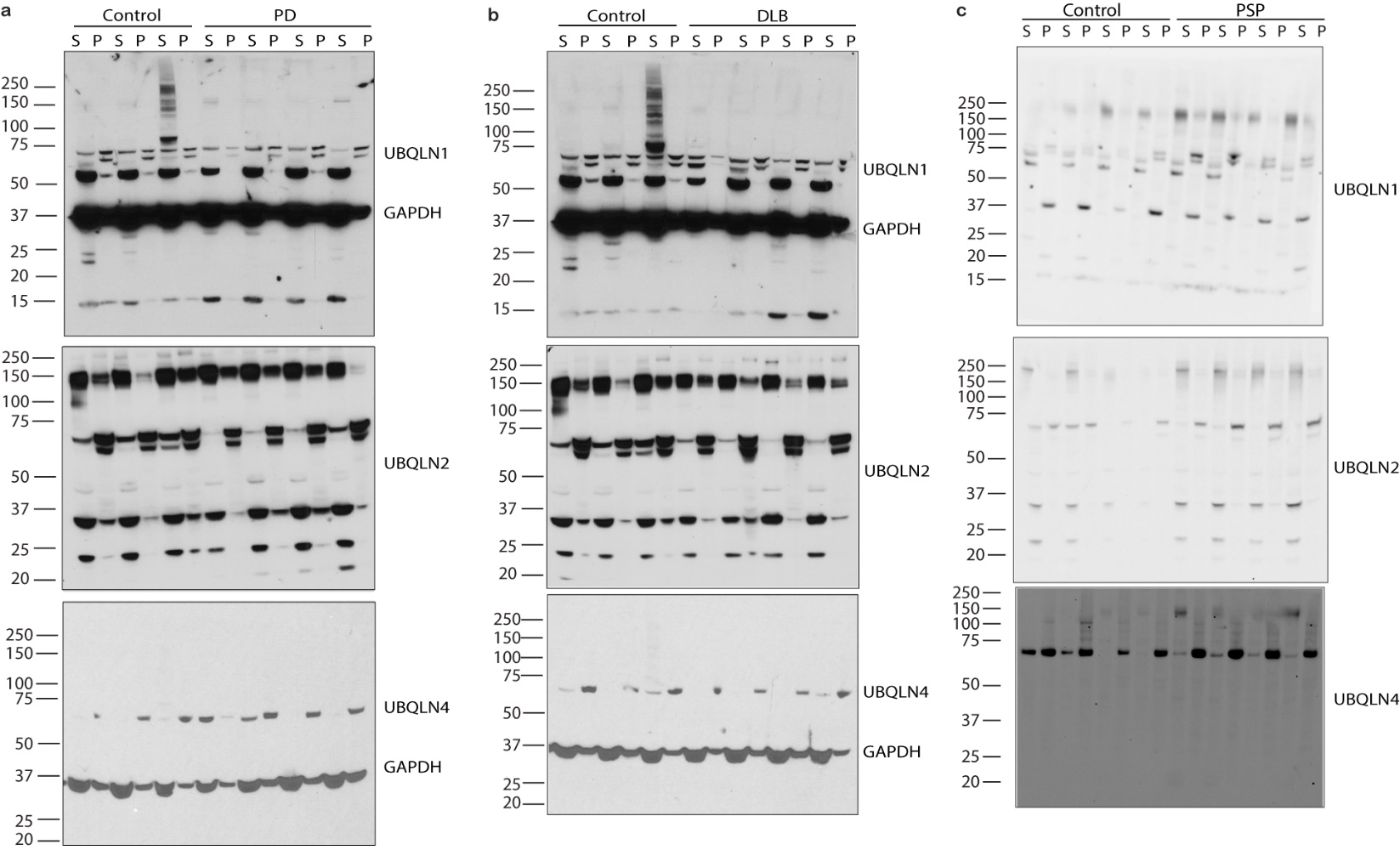

**Supplemental Figure S3.** Full, uncropped Western blots of human brain lysates from Figure 4 detected with UBQLN1, 2 and 4-specific antibodies.

| Human Disease Tissue | | | |
| --- | --- | --- | --- |
| Diagnosis | Sex | Age | PMI* (hrs) |
| Control | Female | 83 | 21 |
| Control | Female | 80 | 19 |
| Control | Male | 100 | 3 |
| Control | Female | 96 | 18 |
| Control | Male | 75 | 9 |
| Control | Male | 65 | 24 |
| Control | Female | 83 | Unknown |
| Control | Male | 71 | 4 |
| Control | Female | 80 | 5 |
| Control | Male | 65 | 14 |
| Control | Female | 74 | 6 |
| Control | Female | 76 | 14 |
| Control | Male | 83 | 28 |
| PD | Male | 78 | 22 |
| PD | Male | 74 | 14 |
| PD | Female | 71 | 7 |
| PD | Male | 86 | 10 |
| PD | Female | 74 | 6 |
| PDD | Male | 81 | 16 |
| DLB | Male | 78 | 12 |
| DLB | Female | 82 | 10 |
| DLB | Male | 84 | 5 |
| DLB | Female | 68 | 24 |
| DLB | Male | 66 | 8 |
| DLB | Male | 86 | Unknown |
| DLB | Male | 66 | 15 |
| DLB | Male | 71 | 5 |
| DLB | Female | 80 | 4 |
| DLB | Female | 57 | 9 |
| DLB | Female | 84 | 6 |
| DLB | Male | 87 | 13 |
| DLB | Male | 72 | 18 |
| DLB | Female | 71 | 12 |
| PSP | Female | 88 | 15 |
| PSP | Male | 64 | 4 |
| PSP | Female | 66 | 12 |
| PSP | Male | 66 | 5 |
| PSP | Male | 73 | 4 |
| PSP | Male | 77 | 12 |
| PSP | Female | 78 | 6 |
| PSP | Male | 73 | 4 |
| PSP | Female | 79 | 6 |
| PSP | Male | 79 | 6 |
| PSP | Male | 73 | 3 |
| PSP | Female | 54 | 12 |

**Supplementary Table** S**1** Human samples used for analysis of UBQLNs. *PMI - postmortem interval.

ANOVA model F statistic and p-value: $F_{3,6}=5.30, p=0.025$

| **Y=UBQLN1 punctum area** | **Coefficients** | **Standard Error** | **t Stat** | **P-value** |
| --- | --- | --- | --- | --- |
| Intercept | 31.8784 | 10.22984 | 3.116216 | 0.00291025 |
| Circularity | -27.0624 | 11.75924 | -2.30137 | 0.025184191 |

ANOVA model F statistic and p-value: $F_{3,6}=8.72, p=0.004$

| **Y=UBQLN2 punctum area** | **Coefficients** | **t Stat** | **P-value** |
| --- | --- | --- | --- |
| Intercept | 18.0250212 | 4.597055818 | 1.39259E-05 |
| Circularity | -13.59759491 | -2.9536822 | 0.004005986 |

ANOVA model F statistic and p-value: $F_{3,6}=4.67, p=0.034$

| **Y=UBQLN4 punctum area** | **Coefficients** | **t Stat** | **P-value** |
| --- | --- | --- | --- |
| Intercept | 22.76455303 | 3.689648536 | 0.000397739 |
| Circularity | -15.95867673 | -2.16149299 | 0.033503036 |

**Supplementary Table S2**. Linear regression analysis of punctum area using circularity as a predictor in the images taken from live HEK293-T cells transfected with the indicated eGFP-labeled ubiquilins.
